# Endogenous opioid system modulates proximal and distal threat signals in the human brain

**DOI:** 10.1101/2024.04.30.591095

**Authors:** Kerttu Seppälä, Vesa Putkinen, Harri Harju, Eleni Rebelos, Jussi Hirvonen, Semi Helin, Johan Rajander, Henry K. Karlsson, Jani Saunavaara, Jukka Hyönä, Lauri Nummenmaa

## Abstract

**BACKGROUND:** Fear promotes rapid detection of threats and appropriate fight-or-flight responses. The endogenous opioid system modulates responses to pain and psychological stressors. Opioid agonists also have also anxiolytic effects. Fear and anxiety constitute major psychological stressors for humans, yet the contribution of the opioid system to acute human fear remains poorly characterized.

**METHODS:** We induced intense unconditioned fear in the subjects by gradually exposing them to a living constrictor snake (threat trials) versus an indoor plant (safety trials). Brain haemodynamic responses were recorded from 33 subjects during functional magnetic resonance imaging (fMRI). In addition, 15 subjects underwent brain positron emission tomography (PET) imaging using [^11^C]carfentanil, a high affinity agonist radioligand for μ-opioid receptors (MORs). PET studies under threat or safety exposure were performed on separate days. Pupillary arousal responses to snake and plant exposure were recorded in 36 subjects. Subjective fear ratings were measured throughout the experiments.

**RESULTS:** Self-reports and pupillometric responses confirmed significant experience of fear and autonomic activation during the threat trials. fMRI data revealed that proximity with the snake robustly engaged brainstem defense circuits as well as thalamus, dorsal attention network, and motor and premotor cortices. These effects were diminished during repeated exposures. PET data revealed that [^11^C]carfentanil binding to MORs was significantly higher during the fear versus safety condition, and the acute haemodynamic responses to threat were dependent on baseline MOR binding in the cingulate gyrus and thalamus. Finally, baseline MOR tone predicted dampening of the haemodynamic threat responses during the experiment.

**CONCLUSIONS:** Preparatory response during acute fear episodes involves a strong motor component in addition to the brainstem responses. These haemodynamic changes are coupled with a deactivation of the opioidergic circuit, highlighting the role of MORs in modulating the human fear response.

## Introduction

Fear acts as a survival intelligence, promoting rapid detection of threats and appropriate fight-or-flight responses by managing information processing priorities and proximity with potentially harmful targets such as predators (Mobbs et al., 2015; Vuilleumier, 2005). The mammalian fear circuit consists of a complex set of midbrain and medial temporal lobe structures, particularly periaqueductal grey, hypothalamus, and amygdala, which interact with prefrontal systems accessing conscious feelings, thus allowing coping with acute and distal threats (Fullana et al., 2016; Tao et al., 2021; Tovote et al., 2015; Zhou et al., 2021). These systems operate flexibly depending on the proximity of the threat. Functional magnetic resonance imaging (fMRI) studies have shown that the imminence of threat switches activation from ventromedial prefrontal cortex (vmPFC) towards periaqueductal grey matter (PAG), reflecting a shift from complex planning of avoidance strategies to automated fight-or-flight response when the predator enters the circa-strike zone (Mobbs et al., 2007; Mobbs et al., 2010). In contrast, activity in the subgenual anterior cingulate cortex is associated with episodes of courage – attempting to regulate the fear and move closer to its target (Nili et al., 2010).

The majority of the imaging studies on the brain basis of human fear have, however, been conducted with biologically unspecific fMRI, and the neuromolecular mechanisms behind acute human fear and its regulation remain elusive. Yet, a bulk of studies suggests that the endogenous μ-opioid receptors system could play a key role in regulating the human fear responses (see review in Meier et al., 2021). Among the three types of opioid receptors (μ, δ, and κ receptors), the μ-receptors (MORs) mediate the effects of endogenous β-endorphins, endomorphins, enkephalins, and exogenous opioid agonists. The predominant action of MORs in the nervous system is inhibitory, but they also have excitatory effects. Multiple OR subtypes are abundantly expressed in the amygdala (Colasanti et al., 2011). MOR expression is also high throughout the human emotion circuit (Nummenmaa & Tuominen, 2018). The MOR system is engaged during various positive and negative emotions (Nummenmaa et al., 2018a; Prossin et al., 2016; Saanijoki et al., 2018; Tuulari et al., 2017), and baseline MOR tone modulates haemodynamic responses to negative affect (Karjalainen et al., 2017; Karjalainen et al., 2018; Sun et al., 2022).

Endogenous MOR tone is negatively associated with subclinical anxiety, suggesting that perturbations in the MOR system might make individuals vulnerable to psychological stressors and subsequent development of clinically relevant anxious symptomatology (Nummenmaa et al., 2020). This is further corroborated by clinical data showing MOR down-regulation in the amygdala and anterior cingulate cortex in PTSD patients exposed to combat versus healthy controls (Liberzon et al., 2007). Relatedly, uncontrolled cohort studies have shown that acute opioid agonist administration following a traumatic event inhibits the development of the post-traumatic stress disorder (PTSD), likely due to inhibition of fear conditioning following the traumatic event (Bryant et al., 2009; Holbrook et al., 2010; Saxe et al., 2001). Finally, pharmacological experiments have established that when exposed to a natural threat (spider), subjects receiving opioid antagonist naloxone kept more distance from the spider, supporting the role of MOR activation in fear suppression (Arntz et al., 1993; Eippert et al., 2008; Kozak et al., 2007; Merluzzi et al., 1991). However, there is currently no direct *in vivo* evidence of the role of the MOR system activation on unconditioned fear in humans, and the contribution of the endogenous MOR system tone on phasic neural responses during acute threat episodes remains unresolved.

### The current study

Here, we tested whether the MOR system is engaged during naturalistic unconditioned threat – exposure to a live snake. We measured endogenous MOR availability with the high-affinity agonist radioligand [^11^C]carfentanil after a neutral baseline state (exposure to a household plant) and after a 10-step modified exposure treatment mimicking clinical exposure therapy for snake phobia (Merluzzi et al., 1991). During a separate fMRI experiment, we repeatedly exposed the subject to the live snake versus the neutral control object (household plant) with varying proximities and modelled the haemodynamic brain responses as a function of subjective fear levels and proximity to the threat. We show that i) acute unconditioned fear engages the endogenous MOR system, ii) this response is paralleled by haemodynamic activity in the midbrain, thalamic and cortical nodes of the human fear circuit, iii) the magnitude of the haemodynamic responses to fear is dependent on the baseline MOR tone, and iv) baseline endogenous MOR tone predicts the suppression of the fear responses during late versus early phases of the experiment.

## Materials and methods

To validate that a live snake would be a potent fearful stimulus, we first conducted an online survey where we asked subjects (n = 786) to report i) how much they are afraid of snakes (1 = not at all, 10 = extremely much) and how much fear and related symptoms they would experience when exposed to a snake or a neutral control object (a household plant) at different proximities. This revealed that, on average, people were moderately strongly afraid of snakes (M = 5.43, SD = 2.80). However, the fear distribution was clearly bimodal with a third of the subjects having moderately low (< 3) and the other third moderately high (> 7) fear of snakes (**Figure S1)**. Both subjective experiences of fear and somatic fear-related sensations were estimated to be higher when the snake was closer to the subject (**Figure S2)**. This confirmed that the live snake exposure would be a valid and potent model for unconditioned fear in the general Finnish population.

### Subjects

The study was approved by the ethics board of the hospital district of Southwest Finland and conducted according to Good Clinical Practice and the Declaration of Helsinki. All subjects signed written informed consent and were informed that they had the right to withdraw at any time without giving a reason. The participants were compensated for their time and travel costs. We recruited only young females since age and sex influence the MOR system function (Kantonen et al., 2020). Females are also, on average, more afraid of snakes than males (Fredrikson et al., 1996; McLean & Anderson, 2009; Tucker & Bond, 1997).

Subjects were recruited via advertisements on email lists, social media, and bulletin boards. Initially, 177 subjects contacted the researchers and were pre-screened for the fear for snakes with a questionnaire based on the short version of SNAQ (Polák et al., 2020; Zsido et al., 2018). This scale contains seven Likert-scale items from SNAQ (scaled from 0 to 100) tapping emotional responses to snakes (e.g. “The way snakes move is repulsive”, “I’m more afraid of snakes than any other animal”). If their average score in the questionnaire was over 70 out of 100, they were deemed eligible for the imaging experiment and were screened for imaging exclusion and inclusion criteria. To maximize the sample size, the lower cut-off for the eye-tracking study was set at 45. Sample sizes for different parts of the study are shown in **Figure S3**. A total of 51 female subjects were studied: 15 in the PET-fMRI experiment (mean age 26.0 years, range 20-42, snake fear scores mean = 79.9, SD = 9.5, Body mass index (BMI) mean 23.2 kg/m^2^, SD = 2.2), 18 for the fMRI experiment (mean age 23.3 years, range 19-27, snake fear scores mean = 75, SD = 13.1, BMI mean 23.6 kg/m^2^, SD = 3.3) and 18 for the eye tracking experiment (mean age 25.7 years, range 20-37, snake fear scores mean = 58.4, SD = 8.1, BMI mean = 23.7 kg/m^2^, SD = 2.5, 2 left-handed). The exclusion criteria included a history of neurological or psychiatric disorders, smoking or use of nicotine products, alcohol and substance abuse, or current use of medication affecting the central nervous system; also breastfeeding or an attempt to become pregnant and the standard MRI and PET exclusion criteria for the subjects in imaging experiments. The study physician screened the subjects for PET imaging for eligibility, and a psychologist screened them for psychiatric disorders with the MINI 6.0 interview (Sheehan et al., 1998). Structural brain abnormalities that are clinically relevant or could bias the analyses were excluded by a consultant neuroradiologist.

### Measuring Autonomic Fear Response with Eye Tracking

Eye movements were measured with Eye Link II system with a 250 Hz sampling rate and a spatial accuracy better than 0.5 degrees of visual angle. Recordings were done in a dimly and constantly lit room. The subjects were seated with the heart rate belt around their chest, the chin on a chin rest, and the hand on a marked place at the table. Next, the eye tracker was set up, calibrated, and validated using a standard 9-point calibration. The experimenter instructed the subject to keep staring at the fixation cross and to concentrate on the emotions that a snake or a plant evoked. It was made evident that the snake or the plant would never actually touch the subjects. Half of the trials involved the snake and half the plant. On each trial, the snake or the plant was brought to the participant’s field of view in either far (screen), moderate (hand), or close (face) proximity from the subject.

Each trial began with a drift correction and recorded spoken instructions (2.2-2.7 s) indicating the type of the trial. Both the experimenter and the subject heard the sounds, and the experimenter moved the object accordingly to the indicated proximity. Subjects were instructed to keep their eyes fixated on the cross shown at the center of the screen throughout the trial. Gaze position and pupil size were measured throughout the 10 s trial, after which the subject reported on a VAS scale 0-10 (0 = not at all, 10 = extremely) their current level of fear. When responding to the VAS scale, the object was out of the subject’s sight. The eye tracker was recalibrated after each of the 12 trials and detrended before each trial. The experiment consisted of 5 sets of 12 snake and 5 sets of 12 plant trials presented in a randomized order. For analysis, subject-wise pupil size time series were cleaned from blinks using an in-house code based on the PhysioData Toolbox (Kret & Sjak-Shie, 2019), baseline corrected (10 ms), and mean pupil sizes were compared between 3 time windows (3-4 s, 5-6 s, 7-8 s) across the conditions (snake vs. plant) and proximities (close, moderate, far).

### PET and MRI measurements

#### Fear Exposure Protocol for PET

Unconditioned fear was induced using a modified version of the fear exposure therapy protocol involving 10 steps (**Table S-1**) with progressively increasing proximity of the snake, leading to potentiation of the fear response. Subjects were first informed about the next step so that they could evaluate if they would be ready to move to the next one. If the subject evaluated the next step as too intense, the current step was repeated. The duration of each step was 60 s. Fear ratings were obtained at 10 seconds and 50 seconds during each step.

### PET Data Acquisition

Subjects underwent two PET scans (threat and safety) in a counterbalanced order. The scans were done at the same time of the day on separate days. Prior to the threat scan, the subjects underwent the 10-step fear exposure protocol (see above), where they were progressively exposed to closer contact with the live snake. To maintain the desired fear level after the exposure and while waiting for the radiotracer injection, the snake was brought repeatedly close to the subject’s lap and hand (approximately 10 minutes). To boost the fear responses, the experimenter had a casual conversation with another researcher during the cannulation, asking for hydrocortisone in case the snake decided to bite. They ensured that the subject heard the conversation. The experimenter placed the hydrocortisone on a research table and wore protective gloves. If a subject asked if the snake was dangerous, the experimenter answered in a following manner: “Don’t worry about it, the snake has bitten only once, and we have the hydrocortisone ready”. Hydrocortisone was not actually needed since the snake was not venomous. When the scanner room was emptied for CT image acquisition, the experimenter sneakily replaced the real snake with a rubber snake and returned the rubber snake in the terrarium for the PET scan. Inside the PET scanner room, the subject could see the rubber snake via an angled mirror, and the snake was kept as close to subject’s head as possible. The rubber snake was used to avoid scatter radiation load to the experimental snake. Only 1/15 subject noticed the sham. During the safety exposure, a routine PET scan was performed, and no external stimulus was presented. Subjects reported their fear levels at the beginning of the experiment and every 10 minutes during imaging.

MOR availability was measured with radioligand [^11^C]carfentanil (Eriksson & Antoni, 2015) synthesized as described previously (Kantonen et al., 2021). Radiochemical purity of the produced [^11^C]carfentanil batches was 98.3 ± 0.42 % (mean ± SD). The injected [^11^C]carfentanil radioactivity was 253 ± 8.96 MBq and molar radioactivity at time of injection 360 ± 240 MBq/nmol corresponding to an injected mass of 0.47 ± 0.42 μg. Subjects were instructed to abstain from smoking and drinking alcohol or caffeine and to avoid physical exercise the day before and the day of the PET scans. The subjects were also told to fast for 3 hours prior to PET imaging. PET imaging was carried out with Discovery 690 PET/CT scanner (GE. Healthcare, US). The tracer was administered as a single bolus via a catheter placed in subject’s antecubital vein, and radioactivity was monitored for 51 minutes. Subject’s head was strapped to the scan table to prevent excessive head movement. T1-weighted MR scans were acquired in a separate session to correct for attenuation and for anatomical reference.

### PET Image Processing and Data Analysis

PET data were preprocessed with the Magia (Karjalainen et al., 2020) toolbox (https://github.com/tkkarjal/magia), an automated neuroimage analysis pipeline developed at the Human Emotion Systems Laboratory, Turku PET Centre. The Magia toolbox runs on MATLAB (The MathWorks, Inc., Natick, MA, USA), and utilizes the methods from SPM12 (www.fil.ion.ucl.ac.uk/spm/) and FreeSurfer (https://surfer.nmr.mgh.harvard.edu/) as well as in-house developed tools for kinetic modeling. PET images were first motion-corrected, and co-registrated to T1-weighted (T1w) MR images, after which T1w image was then processed with Freesurfer for anatomical parcellation. [^11^C]carfentanil uptake was quantified by binding potential relative to non-displaceable binding (*BP*_ND_) in 21 regions (amygdala, caudate, cerebellum, dorsal anterior cingulate cortex, hippocampus, inferior temporal cortex, insula, medulla, midbrain, middle temporal cortex, nucleus accumbens, orbitofrontal cortex, pars opercularis, posterior cingulate cortex, pons, putamen, rostral anterior cingulate cortex, superior frontal gyrus, superior temporal sulcus, temporal pole, and thalamus). *BP*_ND_ was estimated with the simplified reference tissue model (SRTM) in region of interest (ROI (Lammertsma & Hume, 1996) and voxel-level (Gunn et al., 1997) (SRTM) in ROI using the occipital cortex as the reference region. Due to the small sample size, we focused on (SRTM) in region-of-interest ROI analyses, whereas the voxel-level *BP*_ND_ images were used solely for the illustration. Prior to the calculation of voxel-level *BP*_ND_ images, the [^11^C]carfentanil PET images were smoothed using Gaussian kernel to increase signal-to-noise ratio before model fitting (FWHM = 2 mm). *BP*_ND_ images were spatially normalized to MNI152-space and smoothed using a Gaussian kernel (FWHM = 6 mm). Subsequently, regional *BP*_ND_ across the fear and baseline conditions were compared using paired samples t-tests.

### Fear Exposure Protocol for fMRI

During the fMRI experiment, the snake (threat stimulus) and the plant (safety stimulus) were brought to three different proximities (close, moderate, and far distances) from the subject by the experimenter: 1) as close as possible to the subject’s abdomen (close distance) 2) by the subject’s feet (moderate distance) 3) as far as possible on the other side of the room (far distance) by the NordicNeuroLab fMRI screen but still visible for the subject from the gantry via mirror. Each block lasted for 15 seconds and was followed by a VAS rating for fear and a 5 s fixation block during which the experimenter heard via earphones instructions for the next condition. Each condition was repeated 13 times. The Presentation software controlled the timing of the experiment. For logistic reasons, the snake and the plant were presented to the subjects only on one side of the room.

### MRI data acquisition

MR imaging was conducted at Turku University Hospital. The MRI data was acquired using SIGNA, Premier 3T MRI system with SuperG gradient technology (GE Healthcare, Waukesha, WI, USA) and the 48-channel head coil. High-resolution structural images were obtained with a T1w MPRAGE sequence (1 mm^3^ resolution, TR 7.3 ms, TE 3.0 ms, flip angle 8°, 256 mm FOV, 256 × 256 reconstruction matrix). The imaging sequences for the snake and plant scans were identical: 355 functional volumes (15 min 33 s) were acquired with a T2*-weighted echo-planar imaging sequence that is sensitive to the blood-oxygen-level-dependent (BOLD) signal contrast (TR 2600 ms, TE 30 ms, 75° flip angle, 240 mm FOV, 80 × 80 reconstruction matrix, 250 kHz bandwidth, 3.0 mm slice thickness, 45 interleaved axial slices acquired in descending order without gaps).

### MRI data preprocessing, and analysis

The functional imaging data were preprocessed with FMRIPREP (Esteban et al., 2019) (v1.3.0.post2), a Nipype 1.1.8 (Gorgolewski et al., 2011) based tool that internally uses Nilearn 0.5.0 (Abraham et al., 2014). The following preprocessing was performed on the anatomical T1w reference image: correction for intensity non-uniformity, skull-stripping, brain surface reconstruction, spatial normalization to the ICBM 152 Nonlinear Asymmetrical template version 2009c (Fonov et al., 2009) using nonlinear registration with antsRegistration (ANTs 2.2.0) and brain tissue segmentation. The following preprocessing was performed on the functional data: co-registration to the T1w reference, slice-time correction, spatial smoothing with a 6mm Gaussian kernel, automatic removal of motion artifacts using ICA-AROMA (Pruim et al., 2015) and resampling the MNI152NLin2009cAsym standard space. To reveal brain regions encoding the threat value of the snake, the haemodynamic time series during the snake and plant blocks were predicted with trial-wise fear ratings for each subject. To reveal brain regions responding to increased proximity to the threat, we contrasted trials where the snake presented at intermediate proximity was followed by the snake presented at increased (close) versus decreased (far) proximity. To test for the habituation of the fear response, the haemodynamic time series for the fear response was modulated with time (1^st^, 2^nd^, and 3^rd^ third of the experiment), and fear habituation was modelled as positive and negative effects of time. For all analyses, contrast images were generated for positive and negative effects and subjected to the second-level (random effects) model for population-level inference. Statistical threshold was set at 0.05 (FDR corrected at cluster level).

### PET-fMRI fusion analysis

To test whether baseline MOR tone is linked with haemodynamic responses during acute fear, we first extracted mean subject-wise baseline MOR availabilities across the 16 ROIs (see above). Subsequently, the ROI-wise MOR availabilities were correlated with the regional BOLD responses to acute fear characterizing the regional interactions between MOR and BOLD responses during fear. This enabled visualizing the regions where the binding potential estimates most accurate predicted the BOLD responses. A cumulative map of the MOR-dependent BOLD responses was also generated to reveal where the haemodynamic responses are most consistently associated with MOR availability.

## Results

### Self-reports

Subjective fear increased consistently during the exposure trials and the increase was statistically significant (mean slope 0.4, p < .001). Fear remained at a high level until the beginning of the PET scan (mean 2.7) and was significantly higher during the fear versus safety trials throughout the scan (**Figure 1**).

**Figure 1.**
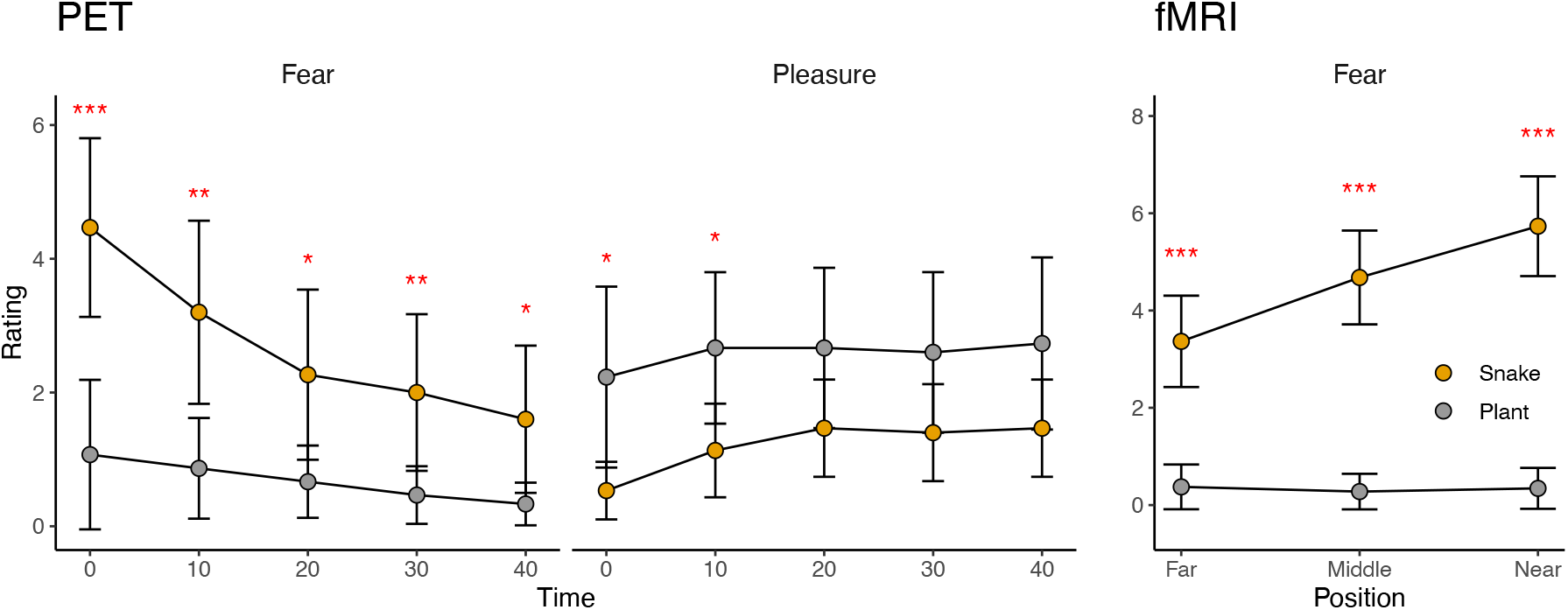
Mean subjective fear and pleasure ratings during the PET scan (left) and mean subjective fear ratings during the fMRI scan (right). Asterisks indicate significant differences between the baseline and snake conditions. *** = p < .001, **= p < .01,* = p < .05.

### Eye tracking

A Condition (Snake, Plant) X Position (Table, Hand, Face) repeated measures ANOVA was run on the mean pupil sizes separately for each time window. The ANOVA and post hoc pair-wise comparison results are shown in **Tables S2-S7**. Eye tracking data (**Figure 2**) revealed that threat exposure led to significant autonomic activity, as indexed by larger pupil sizes when the snake was close to the face or hand versus when it was far away or when the subject was exposed to the plant near the hand or the face at all the analyzed time points (all p < .01) but not when it was on the table (all p > .05). See the full statistics in the **Tables S2-7**.

**Figure 2.**
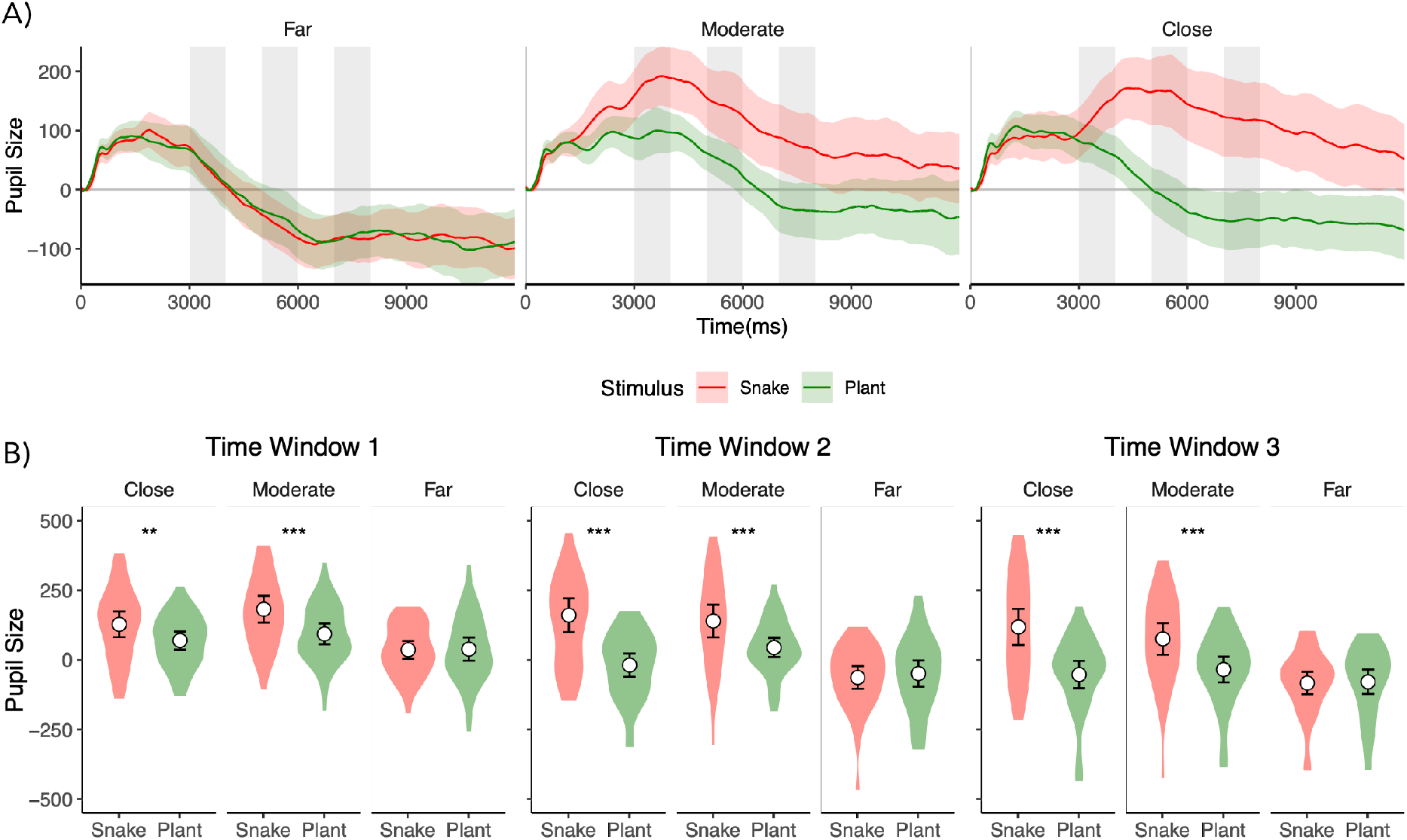
A) Pupillary responses during threat (snake) and safety (plant) trials at the three different distances. Solid lines show mean pupil size, and shaded areas represent 95% CIs. Grey vertical bars indicate the time bins (early (3-4 s), mid (5-6 s), late (7-8s)) where the pupil size was compared across conditions. B) Distributions of mean pupil sizes across the three time-bins for the snake versus plant condition. Asterisks indicate significant differences. ** p < .01, *** p < .001.

### BOLD-fMRI responses associated with fear intensity

A Trial (Snake, Plant) X Position (close, moderate, and far distances) repeated measures ANOVA revealed that mean fear levels were higher (*p* < .001) for in the Snake trials (mean ratings = 4.59, SD = 2.90) compared to the Plant trials (mean rating = 0.33, SD = 1.17) and increased with increasing proximity to the snake but not the plant (Condition X Position interaction: *p* < .001). fMRI data revealed that increasing subjective fear robustly engaged brainstem defense circuits as well as the dorsal attention network and motor and premotor cortices. Significant activations were also observed in the visual cortices. Decreasing fear levels were associated with activation in the amygdala, posterior cingulate cortex, and frontal pole (**Figure 3**).

**Figure 3.**
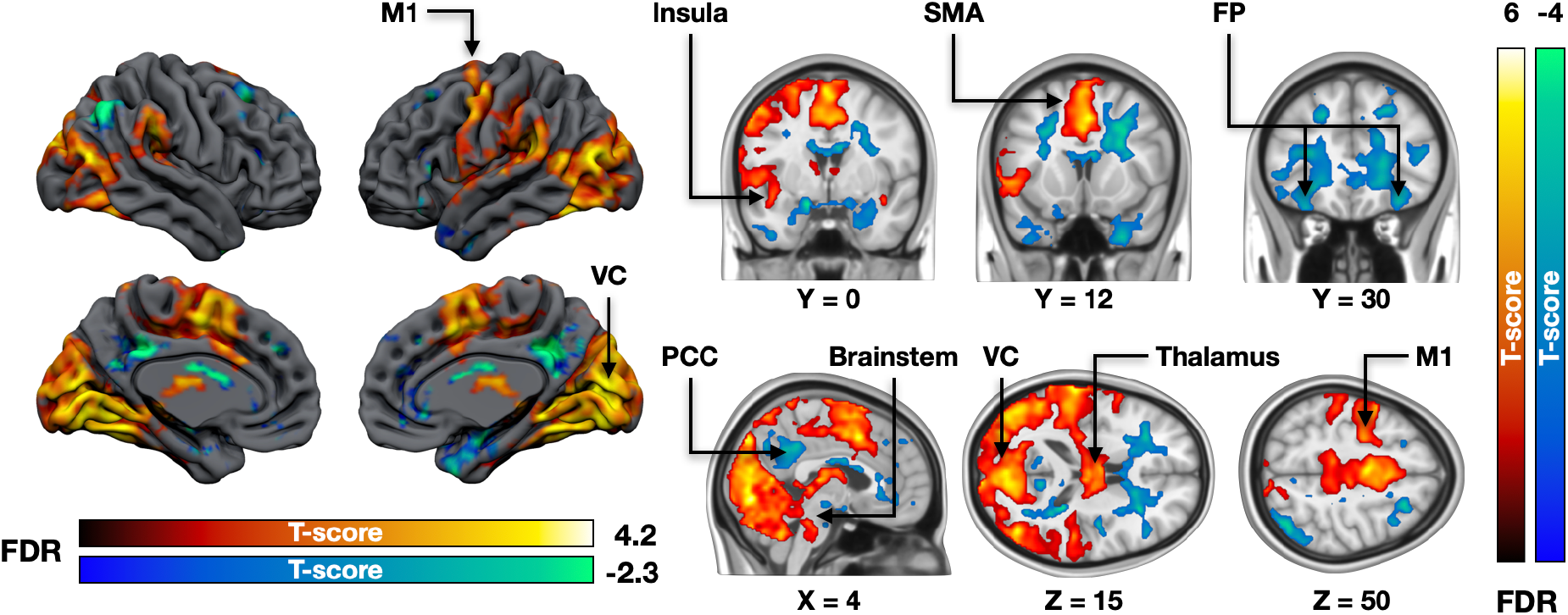
Brain regions responding to increasing (hot colours) and decreasing (cool colours) fear during the experiment. The data are thresholded at p < .05, FDR corrected at cluster level. VC = visual cortex, PCC = posterior cingulate cortex, SMA = supplementary motor area, M1 = primary motor cortex, FP = frontal pole.

### BOLD-fMRI responses associated with threat proximity

Next, we modelled the brain responses associated with increasing versus decreasing proximity of the threat. When the threat stimulus approached the subject, increased activity was observed in the brainstem, cerebellum, insula, and anterior and midcingulate cortices. Additional activations were observed in the primary somatosensory cortex (SI) and inferior frontal and occipital cortices (**Figure 4**). No regions were more active when the threat became more distal.

**Figure 4.**
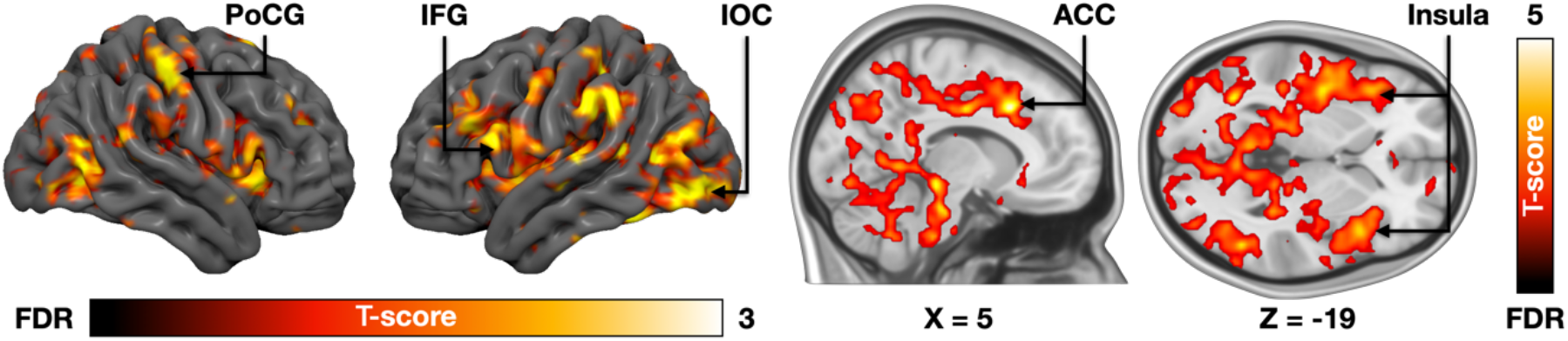
Brain responses to increased proximity of the threat. The data are thresholded at p < .05, FDR corrected at cluster level. ACC = anterior cingulate cortex, IFG = inferior frontal gyrus, IOC = inferior occipital cortex, PoCG = postcentral gyrus.

### Habituation effects for fear in fMRI

We next tested whether the neural responses to fear habituated during the fMRI session. When the fear-dependent haemodynamic responses were modelled as a function of the phase of the experiment (1^st^ / 2^nd^ / 3^rd^), we observed significantly increased responses in the amygdala and brainstem (**Figure 5**). In the opposite contrast, significant effects were observed in the middle cingulate cortex, bilateral fusiform and parahippocampal gyri, left insula and middle temporal cortex.

**Figure 5.**
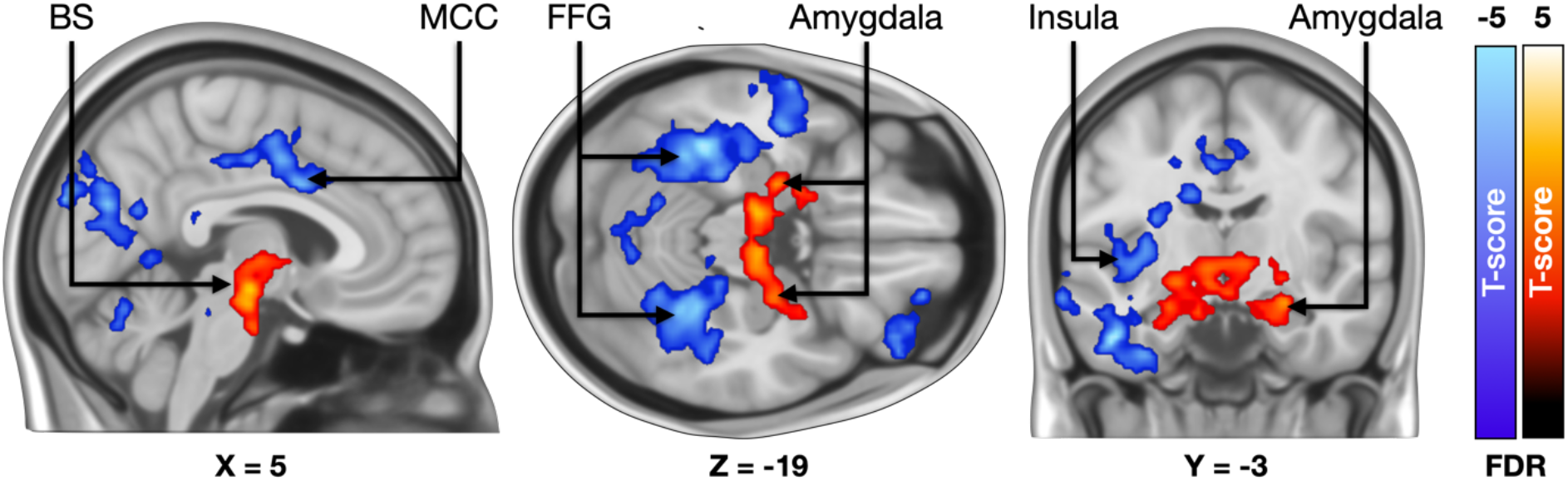
Brain regions where fear-dependent responses became stronger (hot colours) and weaker (blue colours) throughout the experiment. The data are thresholded at p < .05, FDR corrected at cluster level. BS = brainstem, MCC = midcingulate cortex, FFG = fusiform gyri.

### PET

PET data revealed that acute threat influenced the opioid system, as [^11^C]carfentanil *BP*_ND_ was significantly higher during the fear versus safety condition (**Figure 6**). This effect was observed in the left thalamus, hippocampus, middle cingulate cortex, postcentral gyrus, supplementary motor area and precuneus, bilateral middle and superior frontal gyri, and fusiform and lingual gyri. No effects were observed in the opposite contrast.

**Figure 6.**
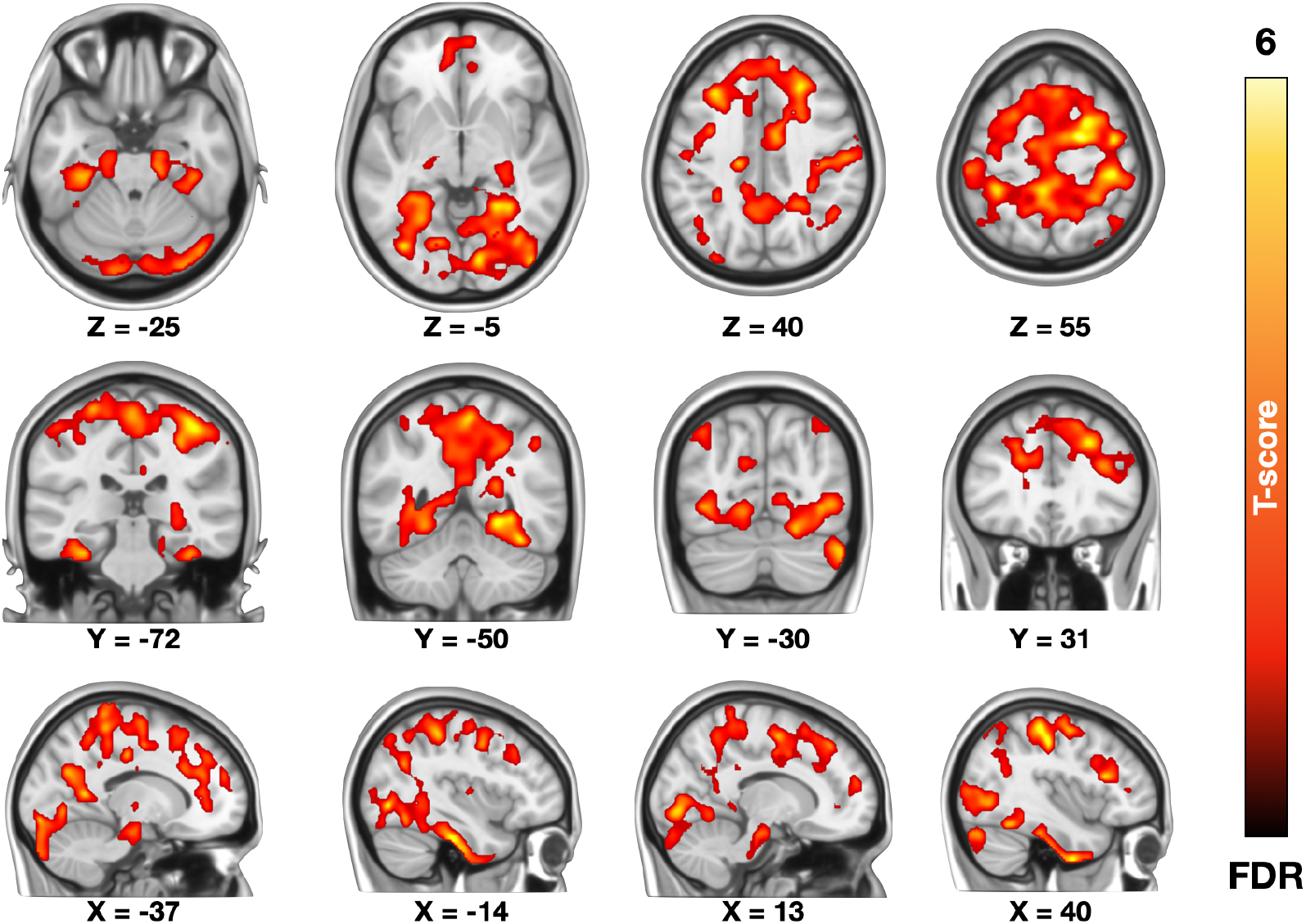
Changes in MOR tone during the fear versus baseline condition; [^11^C]carfentanil *BP*_ND_ was significantly higher during the fear versus safety condition. The data are thresholded at p < .05, FDR corrected at cluster level.

### PET-fMRI fusion analysis

Next, we analyzed the regional interactions between baseline MOR availability and BOLD responses during fear using the PET-fMRI fusion analysis. This revealed a consistent and widespread positive association between MOR tone and fear-dependent haemodynamic responses (**Figure 7**). These effects were observed in limbic and paralimbic regions such as the amygdala and thalamus. Significant effects were also observed in the somatosensory and primary visual cortex and higher-level association areas, brainstem, cerebellum, insula, temporal and frontal cortices. The opposite effects were more limited and observed primarily in anterior portions of cerebellum.

**Figure 7.**
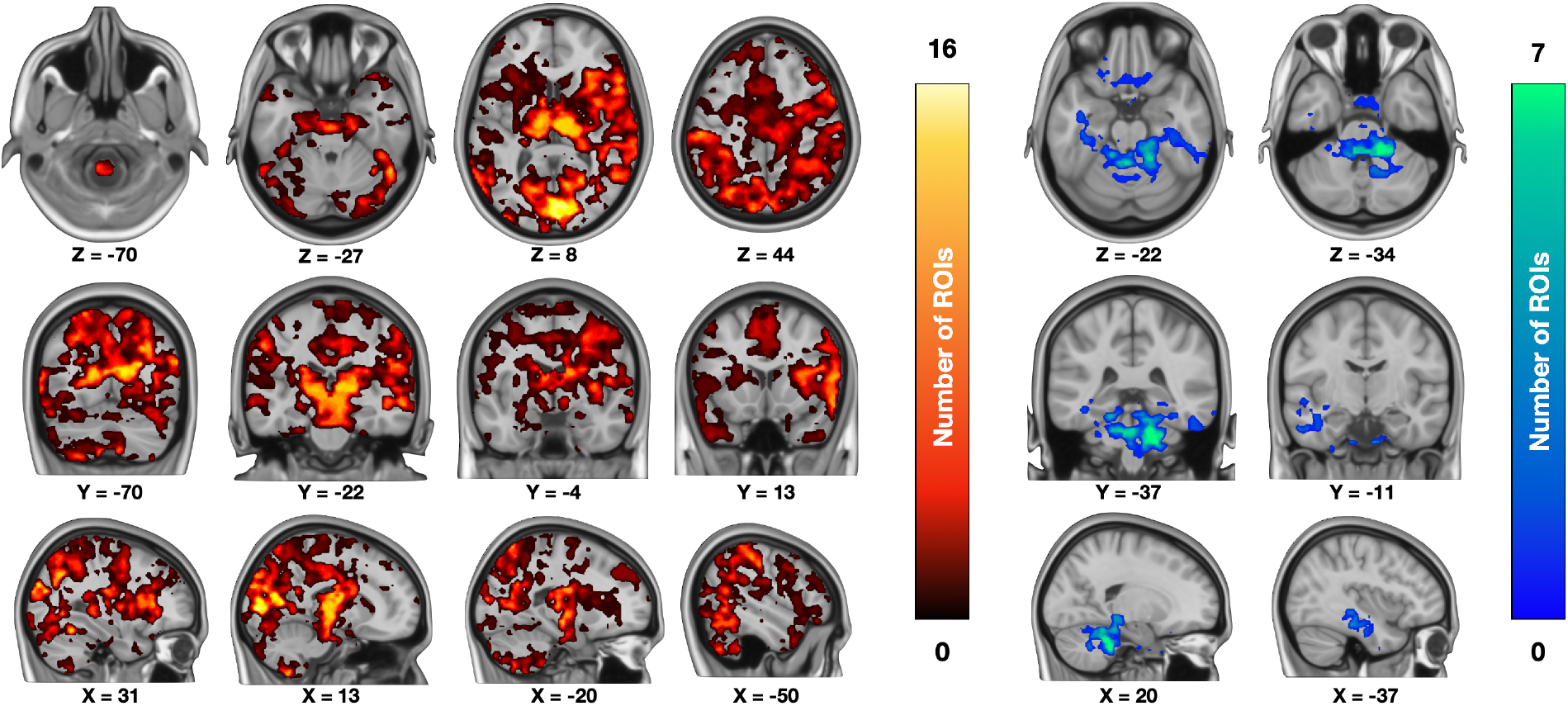
Regional interactions between the MOR system and acute haemodynamic reponses to unconditioned fear. Voxels in this cumulative map show the number of regions (out of 16) whose [^11^C]carfentanil *BP*_ND_ was positively (hot colours) and negatively (cool colors) associated with the subjective fear-dependent BOLD-fMRI responses.

Finally, we tested whether the BOLD response habituation to the fearful stimulus was associated with regional MOR availability. We predicted the strength of the habituation effect (*i*.*e*., changes in BOLD responses during the experiment) with mean regional MOR availabilities. This revealed that the habituation effect was stronger in subjects with higher MOR availability (**Figure 8**). This effect was observed bilaterally in the lateral occipital cortices, lingual gyrus, thalamus, and amygdala. Right-hemispheric activations were also observed in hippocampus, putamen, middle temporal cortices, insula and orbitofrontal cortex and frontal pole.

**Figure 8.**
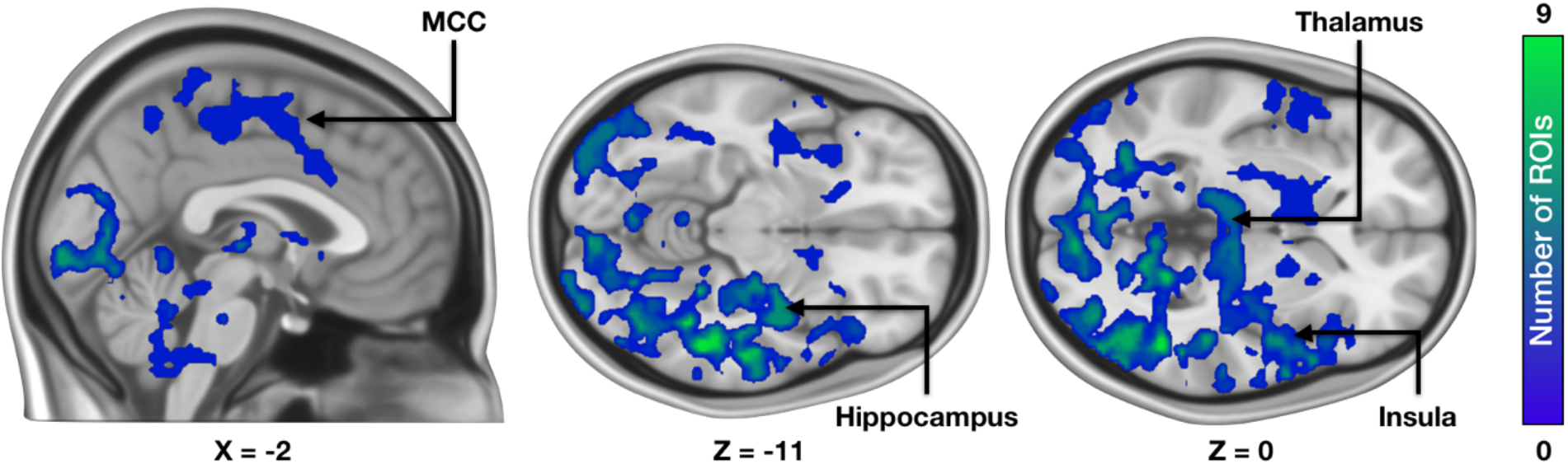
MOR-dependent habituation of haemodynamic fear responses. Voxels in this cumulative map show the number of regions (out of 16) whose [^11^C]carfentanil *BP*_ND_ was positively associated with habituation of the fear-dependent BOLD-fMRI responses throughout the experiment.

## Discussion

Our main finding was that acute fear acutely activated brainstem defense circuits as well as arousal, motor, and attention systems for preparing escape and monitoring survival odds. This acute response was paralleled by autonomic activation indexed by the pupillary responses as well as large-scale deactivation of the endogenous opioidergic system, as indicated by the [^11^C]carfentanil PET data. The MOR system also modulated the acute fear response in that high baseline MOR tone was consistently associated with stronger haemodynamic responses during acute fear. Finally, baseline high MOR tone predicted larger deactivation of cingulate, insular, and hippocampal responses throughout the fMRI experiment. Altogether, these results confirm the role of the endogenous opioid system in modulating acute fear and adapting to threatening situations.

### Haemodynamic responses to acute fear

Acute threat provoked by the proximity of the snake elicited a strong subjective experience of fear, accompanied by autonomic activation as indexed by pupil dilation. On a neural level, the acute fear response was accompanied by increased activity in the brainstem defense circuits and thalamus. These regions manage the acute fight-or-flight responses when the threat was imminent (Mobbs et al., 2007). Significant activations were also observed in the dorsal attention networks and visual cortices. Emotions modulate attentional priorities (Vuilleumier, 2005) and engage the attention circuits consistently across individuals particularly in threatening situations (Nummenmaa et al., 2012). We also found that acute fear led to a significantly increased the activity in the motor and premotor cortices. Emotions prepare the individual for action by adjusting the activation of the cardiovascular, skeletomuscular, neuroendocrine, and autonomic nervous systems (Ekman et al., 1983; Levenson, 2003). Finally, significant effects were observed in the dorsal anterior cingulate cortex, which contributes to the appraisal and expression of negative emotions (Etkin et al., 2011) and integrates lower-order emotional signals into a conscious representation of the emotional state (Saarimäki et al., 2016). Broadly similar effects were observed when haemodynamic responses were modelled as a function of the proximity of the threat, which was expected given that fear levels were consistently higher when the threat stimulus was nearer. Our results thus reveal that the preparatory response during acute fear episodes in fMRI involves a strong motor component in addition to the brainstem responses, indicating an automated preparation of escape behavior.

Although it is widely accepted that the amygdala contributes to emotional processing, vigilance, and relevance detection (Davis & Whalen, 2001; Sander et al., 2003), we did not observe fear or threat proximity-dependent amygdala activity. This occurred despite a well-powered study (n=30), potent and naturalistic fear stimulus triggering consistent autonomic activation, and modelling of the BOLD responses with trial-wise subjective fear ratings. However, a large bulk of neuroimaging studies on human fear has relied on the fear conditioning paradigm. A meta-analysis of such conditioned fear responses has revealed the involvement of SMA/pre-sMA, dACC, anterior insula, ventral striatum, midbrain, and thalamus. Yet, a meta-analysis did not establish significant activations in the amygdala during fear conditioning (Fullana et al., 2016). Recent well-powered fMRI studies have found similar results – a lack of fear-related amygdala activity in the presence of robust peripheral physiological activation but consistent amygdala responses to pictures of human faces (Renée et al., 2021). Accordingly, our data add to the accumulating literature contradicting the notion that the human amygdala is activated during fear, which is demonstrated here under potent naturalistic fear-evoking conditions.

### Opioidergic response to acute fear

Our main finding was that the MOR system responds to acute threat by increasing [^11^C]carfentanil *BP*_ND_, which is traditionally interpreted to occur due to lowered endogenous opioid peptide release leading to increased radioligand binding in the occupancy challenge paradigm (Nummenmaa et al., 2018b). To the best of our knowledge, this is the first *in vivo* demonstration of human MOR system responses during natural, unconditioned fear. We observed widespread modulation of the MOR tone across limbic and paralimbic emotion circuits, as well as somatomotor and frontal cortices. As with fMRI, no effects were observed in the amygdala. This lowered endogenous opioid release is in line with the prior PET studies showing similar opioid system deactivation during negative emotions (Zubieta et al., 2003) as well as large-scale PET studies focusing on individual differences that have deactivation of the MOR system with sustained anxiety (Nummenmaa et al., 2020). Conversely, positive emotions typically lead to increased opioid release in PET studies (Jern et al., 2023; Koepp et al., 2009; Manninen et al., 2017; Tuulari et al., 2017).

All in all, these results show that the endogenous opioid system responds acutely to fear, and individual differences in MOR tone also constitute a molecular factor towards fear and possibly fear-related pathologies. Given the general inhibitory role as well as calming and relaxing functions of μ-receptor agonists (Nummenmaa & Tuominen, 2018), we propose that the presently observed deactivation reflects an increased arousal response during the fight-or-flight situation requiring maximization of physiological and psychological resources for promoting survival. Whereas the MOR system responded to the sustained threat with deactivation, the PET-fMRI fusion analysis revealed that the haemodynamic responses to acute fear were positively associated with baseline MOR availability. In other words, the more MORs the subjects had, the stronger their acute neural fear responses were in limbic and paralimbic regions, including the amygdala and thalamus, as well as cortices and higher-level association areas. This suggests that the MOR system may act at different timescales during threat, with baseline tone associated with acute reactivity and long-term threat exposure leading to sustained MOR downregulation.

### Habituation effects

Self-reports revealed that in our healthy volunteers, even the brief repeated exposure to threat (mimicking exposure therapy for phobia) was sufficient for downregulating self-reported fear of the snake (**Figure 1**). This effect was paralleled by significant changes in the haemodynamic responses to threat in limbic and paralimbic fear circuits, including the brainstem, amygdala, and hippocampus throughout the fMRI experiment. The effect was twofold: responses in the fusiform gyri, hippocampus, insula, and anterior and midcingulate cortices became weaker. Previous fMRI studies have found that activity in the ACC and insula activity is linked with the tendency to withdraw from acute threat (Nili et al., 2010). Thus, the present pattern may indicate a dampening of the escape responses due to fear habituation. In turn, responses in the brainstem and amygdala became stronger. The amygdala effects are noteworthy, as the amygdala did not respond to fear imminence *per se*. However, as its activity was significantly increased throughout the experiment, these results suggest that the amygdala is involved in the adaptation to acute fear in humans. Importantly, the habituation effects in the midcingulate cortex and insula were also dependent on baseline OR availability, suggesting that individual differences in baseline OR tone modulate adaptation to novel threats. This is in line with PET-fMRI studies showing that increased baseline MOR tone buffers against acute haemodynamic responses evoked by negative affect (Karjalainen et al., 2017; Karjalainen et al., 2018; Sun et al., 2022). Additionally, pharmacological studies have found that acute MOR agonist administration effectively inhibits fear learning and the development of PTSD following an acute stressor (Bryant et al., 2009; Holbrook et al., 2010; Saxe et al., 2001). All in all, our results show that the MOR system has an important role in fear regulation and it may act as a buffer against fearful or stressful situations (Nummenmaa et al., 2020) and individual differences in MOR tone may be an important biological mechanism predisposing individuals to sustained fear and anxiety.

### Limitations

Because we scanned only females, we do not know whether our results translate directly to males. The observed *BP*_ND_ changes may reflect altered receptor affinity, internalization, or conformation rather than occupancy by endogenous opioids. Our outcome measure (*BP*_ND_) cannot directly specify which interpretation is most appropriate. As our study was conducted in healthy volunteers, we cannot tell whether the same principles of MOR-dependent fear circuit modulation also occur in subjects with clinical phobia, and this needs to be tested in future studies.

## Conclusions

We conclude that the endogenous opioid system modulates acute fear responses. These effects are observed in i) endogenous MOR system tone changes during threat, as well as in the ii) capacity for the MOR tone to modulate the acute affective and somatomotor threat responses and iii) their downregulation during repeated exposure to threats in the fMRI experiment. Taken together, these results highlight the role of MORs in modulating proximate threats. Clinical studies should further elucidate whether alterations in MOR signaling contribute similarly to clinical phobias and anxiety disorders.

## Supporting information

Seppala et al supplementary material

## Acknowledgements

This study was supported by the Academy of Finland (grants #294897 and #332225), Sigrid Juselius Foundation, Turku University Foundation #080541, Instrumentarium Science Foundation (grants #200061 and #220030), Yrjö Jahnsson foundation #20217451, The Finnish Brain Foundation #20210063, The Päivikki and Sakari Sohlberg Foundation, and State research funding for expert responsibility area (ERVA) of the Turku University Hospital.

## References

Abraham, A., et al. (2014). Machine Learning for Neuroimaging with Scikit-Learn. Frontiers in Neuroinformatics, 8.

Arntz, A., Merckelbach, H., & De Jong, P. (1993). Opioid Antagonist Affects Behavioral Effects of Exposure in Vivo. J Consult Clin Psychol, 61, 865–870.

Bryant, R. A., et al. (2009). A Study of the Protective Function of Acute Morphine Administration on Subsequent Posttraumatic Stress Disorder. Biological psychiatry, 65, 438–440.

Colasanti, A., Rabiner, E. A., Lingford-Hughes, A., & Nutt, D. J. (2011). Opioids and Anxiety. J Psychopharmacol, 25, 1415–1433.

Davis, M., & Whalen, P. J. (2001). The Amygdala: Vigilance and Emotion. Molecular psychiatry, 6, 13–34.

Eippert, F., et al. (2008). Blockade of Endogenous Opioid Neurotransmission Enhances Acquisition of Conditioned Fear in Humans. J Neurosci, 28, 5465–5472.

Ekman, P., Levenson, R. W., & Friesen, W. V. (1983). Autonomic Nervous-System Activity Distinguishes among Emotions. Science, 221, 1208–1210.

Eriksson, O., & Antoni, G. (2015). [11c]Carfentanil Binds Preferentially to Mu-Opioid Receptor Subtype 1 Compared to Subtype 2. Mol Imaging, 14, 476–483.

Esteban, O., et al. (2019). Fmriprep: A Robust Preprocessing Pipeline for Functional Mri. Nat Methods, 16, 111–116.

Etkin, A., Egner, T., & Kalisch, R. (2011). Emotional Processing in Anterior Cingulate and Medial Prefrontal Cortex. Trends Cogn Sci, 15, 85–93.

Fonov, V. S., et al. (2009). Unbiased Nonlinear Average Age-Appropriate Brain Templates from Birth to Adulthood. Neuroimage, 47, S102.

Fredrikson, M., Annas, P., Fischer, H., & Wik, G. (1996). Gender and Age Differences in the Prevalence of Specific Fears and Phobias. Behaviour Research and Therapy, 34, 33–39.

Fullana, M. A., et al. (2016). Neural Signatures of Human Fear Conditioning: An Updated and Extended Meta-Analysis of Fmri Studies. Molecular psychiatry, 21, 500–508.

Gorgolewski, K., et al. (2011). Nipype: A Flexible, Lightweight and Extensible Neuroimaging Data Processing Framework in Python. Frontiers in Neuroinformatics, 5.

Gunn, R. N., Lammertsma, A. A., Hume, S. P., & Cunningham, V. J. (1997). Parametric Imaging of Ligand-Receptor Binding in Pet Using a Simplified Reference Region Model. Neuroimage, 6, 279–287.

Holbrook, T. L., et al. (2010). Morphine Use after Combat Injury in Iraq and Post-Traumatic Stress Disorder. N Engl J Med, 362, 110–117.

Jern, P., et al. (2023). Endogenous Opioid Release Following Orgasm in Man: A Combined Pet-Fmri Study. J Nucl Med.

Kantonen, T., et al. (2020). Interindividual Variability and Lateralization of M-Opioid Receptors in the Human Brain. Neuroimage.

Kantonen, T., et al. (2021). Obesity Risk Is Associated with Altered Cerebral Glucose Metabolism and Decreased Mu-Opioid and Cb1-Receptor Availability International Journal of Obesity.

Karjalainen, T., et al. (2017). Dissociable Roles of Cerebral Mu-Opioid and Type 2 Dopamine Receptors in Vicarious Pain: A Combined Pet-Fmri Study. Cerebral cortex (New York, NY: 1991), 1–10.

Karjalainen, T., et al. (2018). Opioidergic Regulation of Emotional Arousal: A Combined Pet-Fmri Study. Cerebral cortex (New York, NY: 1991).

Karjalainen, T., et al. (2020). Magia: Robust Automated Image Processing and Kinetic Modeling Toolbox for Pet Neuroinformatics. Frontiers in Neuroinformatics, 14.

Koepp, M. J., et al. (2009). Evidence for Endogenous Opioid Release in the Amygdala During Positive Emotion. Neuroimage, 44, 252–256.

Kozak, A. T., et al. (2007). Naltrexone Renders One-Session Exposure Therapy Less Effective: A Controlled Pilot Study. Journal of anxiety disorders, 21, 142–152.

Kret, M. E., & Sjak-Shie, E. E. (2019). Preprocessing Pupil Size Data: Guidelines and Code. Behav Res Methods, 51, 1336–1342.

Lammertsma, A. A., & Hume, S. P. (1996). Simplified Reference Tissue Model for Pet Receptor Studies. Neuroimage, 4, 153–158.

Levenson, R. W. (2003). Blood, Sweat, and Fears - the Autonomic Architecture of Emotion. In P. Ekman, J. J. Campos, R. J. Davidson, & F. B. M. DeWaal (Eds.), Emotions inside Out: 130

Years after Darwin’s the Expression of the Emotions in Man and Animals (Vol. 1000, pp. 348–366). New York: New York Acad Sciences.

Liberzon, I., et al. (2007). Altered Central Micro-Opioid Receptor Binding after Psychological Trauma. Biol Psychiatry, 61, 1030–1038.

Manninen, S., et al. (2017). Social Laughter Triggers Endogenous Opioid Release in Humans. The Journal of Neuroscience, 37, 6125–6131.

Mclean, C. P., & Anderson, E. R. (2009). Brave Men and Timid Women? A Review of the Gender Differences in Fear and Anxiety. Clin Psychol Rev, 29, 496–505.

Meier, I. M., Eikemo, M., & Leknes, S. (2021). The Role of Mu-Opioids for Reward and Threat Processing in Humans: Bridging the Gap from Preclinical to Clinical Opioid Drug Studies. Current Addiction Reports, 8, 306–318.

Merluzzi, T. V., Taylor, C. B., Boltwood, M., & Gotestam, K. G. (1991). Opioid Antagonist Impedes Exposure. J Consult Clin Psychol, 59, 425–430.

Mobbs, D., et al. (2015). The Ecology of Human Fear: Survival Optimization and the Nervous System. Front Neurosci, 9, 22.

Mobbs, D., et al. (2007). When Fear Is Near: Threat Imminence Elicits Prefrontal-Periaqueductal Gray Shifts in Humans. Science, 317, 1079–1083.

Mobbs, D., et al. (2010). Neural Activity Associated with Monitoring the Oscillating Threat Value of a Tarantula. Proc Natl Acad Sci U S A, 107, 20582–20586.

Nili, U., Goldberg, H., Weizman, A., & Dudai, Y. (2010). Fear Thou Not: Activity of Frontal and Temporal Circuits in Moments of Real-Life Courage. Neuron, 66, 949–962.

Nummenmaa, L., et al. (2012). Emotions Promote Social Interaction by Synchronizing Brain Activity across Individuals. Proc Natl Acad Sci U S A, 109, 9599–9604.

Nummenmaa, L., et al. (2020). Lowered Endogenous Mu-Opioid Receptor Availability in Subclinical Depression and Anxiety. Neuropsychopharmacology.

Nummenmaa, L., et al. (2018a). M-Opioid Receptor System Mediates Reward Processing in Humans. Nature Communications.

Nummenmaa, L., Tuominen, L., & Hirvonen, J. (2018b). Simultaneous Pet-Mri Confirms That Cerebral Blood Flow Does Not Confound Pet Neuroreceptor Activation Studies. ACS Chemical Neuroscience, 9, 159–161.

Nummenmaa, L., & Tuominen, L. J. (2018). Opioid System and Human Emotions. Br J Pharmacol, 175, 2737–2749.

Polák, J., Sedláčková, K., Landová, E., & Frynta, D. (2020). Faster Detection of Snake and Spider Phobia: Revisited. Heliyon, 6, e03968.

Prossin, A. R., et al. (2016). Acute Experimental Changes in Mood State Regulate Immune Function in Relation to Central Opioid Neurotransmission: A Model of Human Cns-Peripheral Inflammatory Interaction. Molecular psychiatry, 21, 243–251.

Pruim, R. H. R., Mennes, M., Buitelaar, J. K., & Beckmann, C. F. (2015). Evaluation of Ica-Aroma and Alternative Strategies for Motion Artifact Removal in Resting State Fmri. Neuroimage, 112, 278–287.

Renée, M. V., Joe, B., Scholte, H. S., & Merel, K. (2021). Robust Bold Responses to Faces but Not to Conditioned Threat: Challenging the Amygdala’S Reputation in Human Fear and Extinction Learning. The Journal of Neuroscience, 41, 10278.

Saanijoki, T., et al. (2018). Opioid Release after High-Intensity Interval Training in Healthy Human Subjects. Neuropsychopharmacology.

Saarimäki, H., et al. (2016). Discrete Neural Signatures of Basic Emotions. Cereb Cortex, 6, 2563–2573.

Sander, D., Grafman, J., & Zalla, T. (2003). The Human Amygdala: An Evolved System for Relevance Detection. Rev Neurosci, 14, 303–316.

Saxe, G., et al. (2001). Relationship between Acute Morphine and the Course of Ptsd in Children with Burns. J Am Acad Child Adolesc Psychiatr, 40, 915–921.

Sheehan, D. V., et al. (1998). The Mini-International Neuropsychiatric Interview (M.I.N.I.): The Development and Validation of a Structured Diagnostic Psychiatric Interview for Dsm-Iv and Icd-10. J Clin Psychiatry, 59 Suppl 20, 22-33;quiz 34-57.

Sun, L., et al. (2022). Mu-Opioid Receptor System Modulates Responses to Vocal Bonding and Distress Signals in Humans. Phil Trans B.

Tao, D., et al. (2021). Where Does Fear Originate in the Brain? A Coordinate-Based Meta-Analysis of Explicit and Implicit Fear Processing. Neuroimage, 227, 117686.

Tovote, P., Fadok, J. P., & Lüthi, A. (2015). Neuronal Circuits for Fear and Anxiety. Nature Reviews Neuroscience, 16, 317–331.

Tucker, M., & Bond, N. W. (1997). The Roles of Gender, Sex Role, and Disgust in Fear of Animals. Personality and Individual Differences, 22, 135–138.

Tuulari, J. J., et al. (2017). Feeding Releases Endogenous Opioids in Humans. J Neurosci, 37, 8284–8291.

Vuilleumier, P. (2005). How Brains Beware: Neural Mechanisms of Emotional Attention. Trends Cogn Sci, 9, 585–594.

Zhou, F., et al. (2021). A Distributed Fmri-Based Signature for the Subjective Experience of Fear. Nat Commun, 12, 6643.

Zsido, A. N., et al. (2018). Short Versions of Two Specific Phobia Measures: The Snake and the Spider Questionnaires. Journal of anxiety disorders, 54, 11–16.

Zubieta, J. K., et al. (2003). Regulation of Human Affective Responses by Anterior Cingulate and Limbic Mu-Opioid Neurotransmission. Arch Gen Psychiatry, 60, 1145–1153.

