## Supplementary material for "Endogenous opioid system modulates proximal and distal threat signals in the human brain": Seppala et al supplementary material

### Self-reports for snake fear

We conducted an online survey (n= 786) using Google Docs to estimate the typical characteristics of fear for snakes in the population. The link to the questionnaire was posted to social media, various email lists and bulletin boards. Subjects were asked to evaluate on a scale from 1 to 10 (1 = not at all, 10 = extremely much) how much they fear snakes. Additionally, subjects were asked to evaluate (1 = not at all, 10 = extremely much) the intensity of fear, disgust, and fear-related somatic sensations (sweating and heart racing) when they would encounter a snake at different proximities ranging from looking at a picture of a snake to looking at a snake crawling on one's bare abdomen. The distribution for the fear for snakes is shown in **Figure S-1**. On average, people are moderately scared of snakes (M = 5.43, SD = 2.80), however, the fear distribution is clearly bimodal, where a third of subjects have moderately low (<3) and other third moderately high (>7) fear of snakes. The predicted fear, disgust, heart rate, and sweating increase when the snake is imagined to be closer to the subject. As expected, the strongest affective and somatic sensations are observed when the snake is imagined to be on the subjects' abdomen (**Figure S-2**).

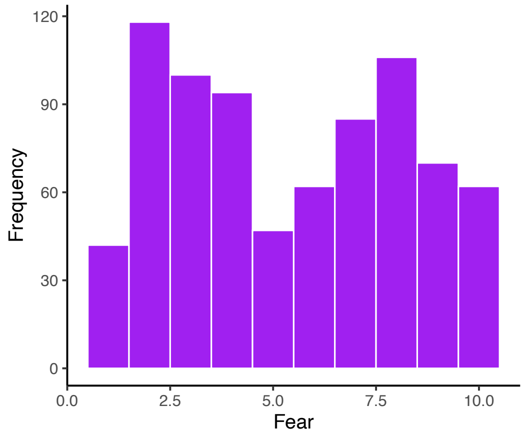


**Figure S-1.** Bimodal distribution of fear for snakes on a scale from 1 (not at all) to 10 (extremely much) in the sample (n=786) of Finnish population.

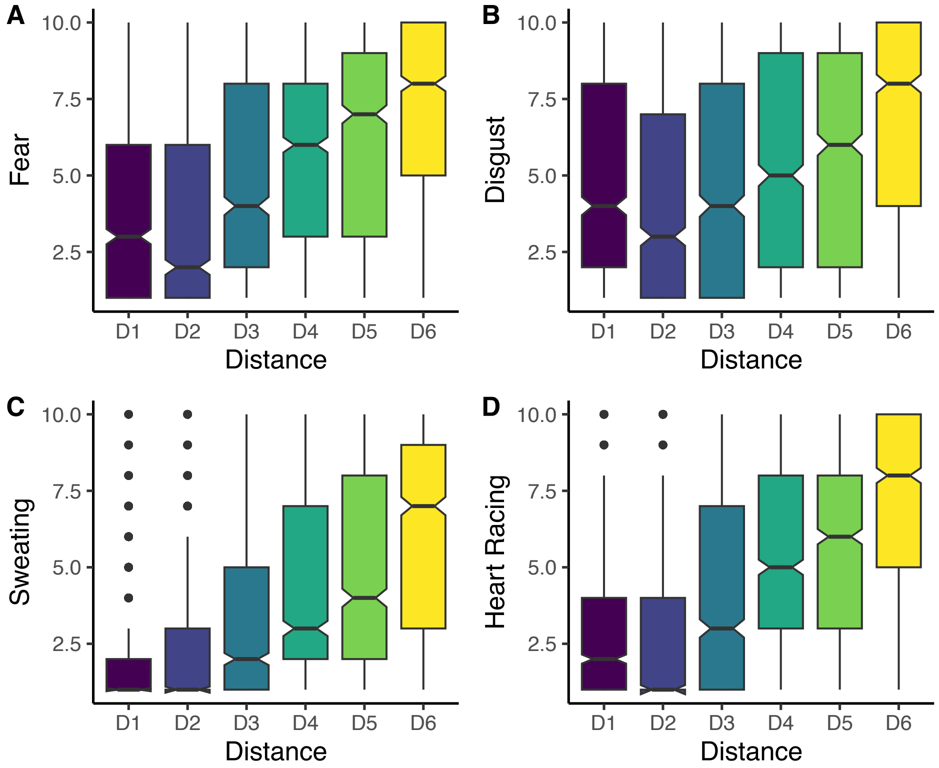


Figure S-2. Distribution of the estimated ratings for fear, disgust, sweating, and heart rate with different imagined proximities of a snake: D1 = looking at a picture of a snake, D2 = looking at a snake in a closed transparent container, D3 = holding a closed, transparent container with a snake close to one's face, D4 = touching a snake with a hand while wearing thick gloves and a long-sleeved shirt, D5 = looking at a snake crawling freely on the floor, D6 = letting a snake crawl on one's bare abdomen.

### Study outline

The main branch of the subjects involved 15 subjects undergoing both PET and fMRI studies. To limit radiation load, only 15 subjects were studied with PET and an additional 15 subjects were recruited for the fMRI study. Due to the already demanding PET protocol, the eye tracking validation protocol was performed only in the subjects who participated in the fMRI studies and in 18 additional subjects.

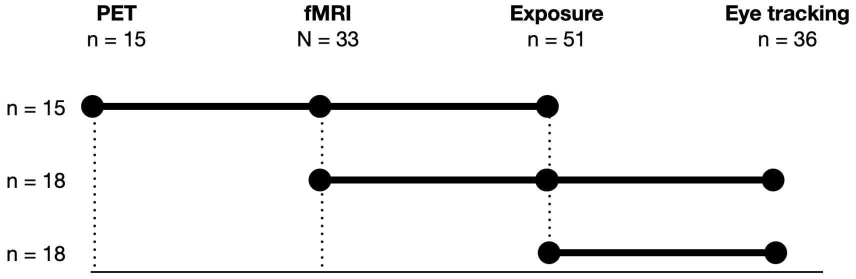


Figure S-3. Overview of the experimental design and the number of subjects in each branch. Note that the figure does not reflect the order of the tasks for each subject.

**Table S-1.** Steps used in the fear exposure protocol.

| Step | Contents |
|---|---|
| 1. | Baseline: subject sits quietly in a chair. |
| 2. | Subject looks at a picture of a snake. |
| 3. | Subject holds a picture of the snake. |
| 4. | Snake is held at 3 m from the subject. |
| 5. | Snake is held at 1.5 m from the subject. |
| 6. | Snake is held at 0.5 m from the subject. |
| 7. | Subject touches the container with a finger. |
| 8. | Subject touches the container with both hands. |
| 9. | Experimenter removes the lid of the container. |
| 10. | Experimenter holds the snake close to the subject's face. |

**Table S-2.** Main effects and interactions for time window 1

| Effect | DFn | DFd | SSn | SSd | F | p | p<.05 | ges |
|---|---|---|---|---|---|---|---|---|
| Intercept | 1 | 34 | 1790404.45 | 1726330.6 | 35.261931 | 0.0000010 | * | 0.3916845 |
| Condition | 1 | 34 | 121291.39 | 299357.0 | 13.775886 | 0.0007339 | * | 0.0417969 |
| Position | 2 | 68 | 356950.43 | 368369.2 | 32.946062 | 0.0000000 | * | 0.1137660 |
| Condition X Position | 2 | 68 | 75676.09 | 386576.1 | 6.655835 | 0.0022914 | * | 0.0264944 |

**Table S-3.** Contrast tests between plant and snake at different positions for time window 1

| Contrast | Position | Estimate | SE | df | T ratio | p-value |
|---|---|---|---|---|---|---|
| Plant - Snake | Face | -58.082534 | 19.60297 | 97.34463 | -2.962946 | 0.0038300 |
| Plant - Snake | Hand | -88.740127 | 19.60297 | 97.34463 | -4.526873 | 0.0000170 |
| Plant - Snake | Table | 2.625542 | 19.60297 | 97.34463 | 0.133936 | 0.8937297 |

**Table S-4.** Main effects and interactions for time window 1

| Effect | DFn | DFd | SSn | SSd | F | p | p<.05 | ges |
|---|---|---|---|---|---|---|---|---|
| Intercept | 1 | 34 | 281591.4 | 2009713.2 | 4.763917 | 0.0360648 | * | 0.0656220 |
| Condition | 1 | 34 | 396089.9 | 513418.5 | 26.230177 | 0.0000119 | * | 0.0899058 |
| Position | 2 | 68 | 902122.3 | 822511.3 | 37.290867 | 0.0000000 | * | 0.1836701 |
| Condition X Position | 2 | 68 | 330107.5 | 663878.9 | 16.906179 | 0.0000011 | * | 0.0760681 |

**Table S-5.** Contrast tests between plant and snake at different positions for time window 1

| Contrast | Position | Estimate | SE | df | T ratio | p-value |
|---|---|---|---|---|---|---|
| Plant - Snake | Face | -179.41989 | 25.68171 | 97.3725 | -6.9862899 | 0.0000000 |

| Contrast | Position | Estimate | SE | df | T ratio | p-value |
|---|---|---|---|---|---|---|
| Plant - Snake | Hand | -95.40694 | 25.68171 | 97.3725 | -3.7149758 | 0.0003391 |
| Plant - Snake | Table | 14.24838 | 25.68171 | 97.3725 | 0.5548063 | 0.5802993 |

**Table S-6.** Main effects and interactions for time window 1

| Effect | DFn | DFd | SSn | SSd | F | p | p<.05 | ges |
|---|---|---|---|---|---|---|---|---|
| Intercept | 1 | 34 | 14447.94 | 2641045.0 | 0.1859983 | 0.6689906 | | 0.0032051 |
| Condition | 1 | 34 | 446783.72 | 427068.9 | 35.5695467 | 0.0000010 | * | 0.0904408 |
| Position | 2 | 68 | 546100.66 | 678822.9 | 27.3523799 | 0.0000000 | * | 0.1083665 |
| Condition X Position | 2 | 68 | 277400.97 | 746349.8 | 12.6370139 | 0.0000215 | * | 0.0581470 |

**Table S-7.** Contrast tests between plant and snake at different positions for time window 1

| Contrast | Position | Estimate | SE | df | T ratio | p-value |
|---|---|---|---|---|---|---|
| Plant - Snake | Face | -171.060365 | 25.63937 | 101.5715 | -6.6717844 | 0.0000000 |
| Plant - Snake | Hand | -110.007738 | 25.63937 | 101.5715 | -4.2905784 | 0.0000407 |
| Plant - Snake | Table | 4.316419 | 25.63937 | 101.5715 | 0.1683512 | 0.8666418 |
